# Magnified interaural level differences enhance binaural unmasking in bilateral cochlear implant users

**DOI:** 10.1101/2024.06.03.597254

**Authors:** Benjamin N. Richardson, Jana M. Kainerstorfer, Barbara G. Shinn-Cunningham, Christopher A. Brown

## Abstract

Bilateral cochlear implant (BiCI) usage makes binaural benefits a possibility for implant users. Yet for BiCI users, limited access to interaural time difference (ITD) cues and reduced saliency of interaural level difference (ILD) cues restricts perceptual benefits of spatially separating a target from masker sounds. The present study explored whether magnifying ILD cues improves intelligibility of masked speech for BiCI listeners in a “symmetrical-masker” configuration, which ensures that neither ear benefits from a long-term positive target-to-masker ratio (TMR) due to naturally occurring ILD cues. ILD magnification estimates moment-to-moment ITDs in octave-wide frequency bands, and applies corresponding ILDs to the target-masker mixtures reaching the two ears at each specific time and frequency band. ILD magnification significantly improved intelligibility in two experiments: one with NH listeners using vocoded stimuli and one with BiCI users. BiCI listeners showed no benefit of spatial separation between target and maskers with natural ILDs, even for the largest target-masker separation. Because ILD magnification relies on and manipulates only the mixed signals at each ear, the strategy never alters the monaural TMR in either ear at any time. Thus, the observed improvements to masked speech intelligibility come from binaural effects, likely from increased perceptual separation of the competing sources.

## I. INTRODUCTION

Bilateral cochlear implant (BiCI) use has become more common in recent years, in part because of the potential to provide CI users access to binaural cues. This can help address poor spatial hearing outcomes in BiCI users, including degraded performance on sound localization tasks (Dorman et al., 2016; Grantham et al., 2008; Jones et al., 2014) and spatial release from masking (SRM) tasks (D’Onofrio et al., 2020; Loizou et al., 2009) compared to their normal-hearing peers. These outcomes are due in part to poor sensitivity to interaural time differences (Noel and Eddington, 2013).

Historically, CIs have not provided a very robust representation of acoustic inputs (Lorenzi et al., 2006; Moore, 2008), especially interaural differences important in spatial hearing (Anderson et al., 2024; Grantham et al., 2008; Laback et al., 2015). Specifically, traditional coding strategies do not represent temporal fine structure (TFS); moreover, because processors are not synchronized across the ears, even interaural envelope differences are not very reliable. Several CI stimulation strategies have attempted to re-introduce or extend interaural time difference (ITD) information in BiCIs (van Hoesel et al., 2009; Long et al., 2003; Srinivasan et al., 2020; Thakkar et al., 2023), or TFS more generally (Ausili et al., 2020). However, there are other factors that reduce perceptual sensitivity, including current spread, frequency limitations on phase locking, and interaural asymmetry in electrode position (Anderson et al., 2024; Gray et al., 2021; Laback et al., 2015). In addition, reduced TFS sensitivity arising from the impaired auditory systems of CI users limits spatial hearing benefits even when devices are linked (Kan et al., 2015).

SRM can arise from monaural TMR improvements at the ear nearer the target, as well as “binaural unmasking” effects based on interaural computations. In normal hearing (NH) listeners, interaural level differences (ILDs) can provide benefits to speech understanding both by affecting the monaural mixtures reaching each ear and by changing the binaural comparisons available to the listener. BiCI users retain some sensitivity to interaural level differences (Grantham et al., 2008), and thus may achieve SRM from salient ILDs through both monaural and possibly binaural effects, if the ILDs are sufficiently large.

Monaural aspects of SRM come about from acoustic changes in the mixture of target and masker signals reaching an ear. When two competing sound sources arrive at a listener’s head from different directions, the target sound is closer to one ear than is the masker, resulting in a greater acoustic target-to-masker energy ratio (TMR) in that ear, a source of benefit termed the better-ear effect (Glyde et al., 2013a, 2013b). Even in situations where there is no long-term average TMR benefit, ILDs can provide dynamic short term better-ear advantages (“glimpses”) at moments when one of the maskers has an energy “dip” (Bronkhorst and Plomp, 1992; Gibbs et al., 2022; Glyde et al., 2013a, 2013b).

Spatial benefits that result from the comparison across both ears, beyond monaural effects, are referred to as binaural unmasking (Culling and Lavandier, 2021; Dieudonné and Francart, 2019). Perceived spatial separation can provide binaural unmasking (Freyman et al., 1999). While ITD cues often are thought to dominate binaural unmasking effects, ILDs can also contribute to the perception of spatial differences in competing sources, thereby promoting sound segregation and supporting selective attention based on perceived source laterality (Best et al., 2005; Ihlefeld and Shinn-Cunningham, 2008; Middlebrooks and Waters, 2020)producing SRM.

Naturally occurring ILDs thus support selective attention in NH listeners when target and masker sound sources are sufficiently spatially separated through both long-term and dynamic better-ear effects, and also by promoting sound source segregation and enabling spatial selective attention. How ILD information can help BiCI users perform selective attention is an open question. Here, we are interested in whether enhancing ILDs can enhance binaural unmasking in BiCI listeners in spatial configurations where monaural better-ear effects are minimized.

BiCI configurations retain some ILD information, which means that listeners using such devices can benefit from a positive TMR at the ear nearer the target (Litovsky et al., 2009; Loizou et al., 2009). However, ILDs are reduced after CI processing (Dorman et al., 2014; Gray et al., 2021), due to various factors, including dynamic range compression associated with frontend automatic gain control that operates independently at the two ears (Archer-Boyd and Carlyon, 2019, 2021; Spencer et al., 2019), backend mapping (Khing et al., 2013), and independent peak picking (Gray et al., 2021). These limitations on the perceptibility of better-ear effects help explain why bilateral CI users show both reduced glimpsing (Gibbs et al., 2022; Hu et al., 2018) and a reduced ability to selectively attend to a target in the presence of a spatially separated masker (Akbarzadeh et al., 2020; Goupell et al., 2016). As a result, natural ILD cues alone are not sufficient to support SRM in BiCI users (Ihlefeld and Litovsky, 2012).

Magnifying low-frequency ILD information (i.e., increasing ILD cue magnitudes to be larger than those that occurs naturally) can significantly improve speech intelligibility in BiCI users compared to presenting naturally occurring, unprocessed ILDs (Brown, 2014). Specifically, for a target talker to the left of midline and a masker talker to the right, magnification of ILDs increases SRM (Brown, 2014). Because ILD magnification works on the target-masker mixtures at each ear (changing both target and masker identically in an ear), it does not change the TMR in either ear. This suggests that the benefits from ILD magnification may be due to improved spatial selection of the target stream. Although ILD magnification would require synchronization across devices, ILD cues are simple to introduce into existing BiCI strategies because they only require sensitivity to differences in stimulation intensity at each ear, which is better preserved in electric hearing than is timing information.

The current study extends these previous results by comparing different ILD magnification schemes. We tested speech intelligibility in both NH listeners presented with vocoded stimuli (Experiment 1) and in BiCI users (Experiment 2). We used three ILD magnification conditions: No magnification, which used naturally occurring spatial cues via non-individualized head-related transfer functions (HRTFs; (Gardner and Martin, 1994); Low-Frequency magnification, which applied the same HRTF-based natural cues, then applied ILD magnification in frequency bands below 2 kHz where naturally-occurring ILDs are less robust, similar to Brown (Brown, 2014); and Broadband magnification, which applied ILD magnification in frequency bands spread across a wider range of frequencies (125-5500 Hz). Crucially, the Broadband magnification approach extends magnified ILDs into mid-frequencies (i.e. 1-5 kHz), which are important for speech understanding (DePaolis et al., 1996).

We also changed the number of processing bands used, compared to what was used earlier (Brown, 2014). Though listeners benefit from magnification applied to 20 ERB-wide bands below 2 kHz (Brown, 2014), we do not know how results would change if magnification was applied to fewer independent frequency bands. This is an important practical consideration if the strategy is to be implemented in BiCIs, since fewer frequency bands reduces processor cycles, which in turn reduces processing latency and improves battery life. The current study does not parametrically vary processing band number due to testing time constraints. As few as six one-octave processing bands significantly improved localization performance (Brown, 2018), so here we explored whether benefits could be obtained using only four bands.

The current study also builds on previous iterations of ILD magnification by testing performance using symmetrical-masker spatial configurations. We employed these spatial configurations for a number of reasons. First, we wanted to observe the effectiveness of the strategy in a more acoustically complex scene than the single-masker configuration used previously (Brown, 2014). We also wanted to control for better-ear acoustic benefits that arise from naturally-occurring ILD cues in asymmetrical target-masker configurations, which lead to improved long-term monaural TMR (Glyde et al., 2013a) and allow many opportunities for shorter-term glimpsing (Bronkhorst and Plomp, 1992; Gibbs et al., 2022). In a symmetrical-masker configuration with the target at midline and a masker on either side, better-ear benefits do not increase as target-masker separation increases; instead, each masker moves closer to its ipsilateral ear while target position does not change. Shorter-term glimpsing will also be reduced compared to asymmetrical configurations. Given how we implement ILD magnification, monaural TMR improvements cannot explain any of the perceptual benefits we observe; the algorithm is applied to the mixed signals at each ear, decreasing the overall level of one ear’s total acoustic input (depending on the dominant ITD at that time and frequency). Given this, the monaural TMRs in the two ears do not change with magnification: one ear’s signal will be unchanged, and the other’s overall level will decrease without changing the TMR.

In NH listeners with vocoded stimuli, we hypothesized that naturally occurring ILDs alone would be insufficient to produce SRM at small target-masker separations, but would be sufficient at large separations where ILDs are larger. We further hypothesized that ILD magnification would improve task performance by increasing the perceived spatial separation of target and maskers, allowing more successful spatial selective attention. We expected Low-frequency magnification to yield better performance than naturally occurring ILDs and that Broadband magnification would produce the best performance (and the strongest SRM).

For BiCI listeners, we hypothesized that naturally occurring ILDs would fail to produce any SRM (Brown, 2014; Ihlefeld and Litovsky, 2012), even for large target-masker separations. We hypothesized that Low-frequency ILD magnification would improve performance on our SRM task, and that Broadband ILD magnification would produce even better performance.

## II. MATERIALS AND METHODS

### A. Participants

Sixteen NH listeners (Experiment 1) and seven BiCI users (Experiment 2) participated. A hearing screening with NH listeners confirmed that all listeners had audiometric thresholds of 25 dB HL or lower at octave frequencies between 125 and 8000 Hz. BiCI users had at least 12 months of experience with both devices. During testing, they used their everyday program without modification, except for a minor volume adjustment to ensure that sounds were at a comfortable overall level. The NH group were all between the ages of 19 and 37. The ages and devices used by each BiCI user are shown in Table I. All participants were compensated for their participation either monetarily or with course credit. All procedures were approved by the Institutional Review Board of the University of Pittsburgh.

**TABLE I.** Age and device used in each ear, for each BiCI user.

| Subject # | Age (years) | Left Ear Device | Right Ear Device |
| --- | --- | --- | --- |
| 1 | 68 | Cochlear Nucleus 5 | Cochlear Nucleus 5 |
| 2 | 72 | Cochlear Nucleus 5 | Cochlear Nucleus 5 |
| 3 | 76 | Cochlear Nucleus 5 | Cochlear Nucleus 5 |
| 4 | 75 | MED-EL Opus 2 | MED-EL Opus 2 |
| 5 | 64 | Cochlear Nucleus 5 | Cochlear Nucleus 5 |
| 6 | 52 | AB Harmony | AB Harmony |
| 7 | 72 | Cochlear Nucleus 5 | Cochlear Nucleus 5 |

### B. Stimuli and Signal Processing

Target speech was drawn from the CUNY sentence corpus (Boothroyd et al., 1985) produced by a female talker. Masker stimuli were drawn from the IEEE corpus (Rothauser, 1969), which were produced by a different female talker. Masker sentences were concatenated if needed and then truncated to be equal in duration to the target sentence chosen for a given trial.

Stimulus processing and presentation, as well as experiment control, were accomplished using the Python programming language. All signal processing was performed offline (see schematization in Fig. 1). To generate stimuli, each of the two masker sentences were first adjusted in level relative to the target sentence level to achieve a TMR of +2 dB. Then, naturally occurring binaural cues were applied separately to the target and maskers by convolving each with appropriate non-individualized HRTFs ((Gardner and Martin, 1994). Target sentences were simulated at midline, while the maskers were symmetrically positioned around midline at azimuths of ± 0, 15, 30, 45, 60, 75, or 90 degrees, all at ear height (0 degrees elevation). Following HRTF spatialization, target and masker tokens were summed into left and right ear channels. Left and right ear channels were bandpass filtered between 125 and 5500 Hz to simulate the limited bandwidth of CI processing and to set the effective bandwidth of the signals to be comparable to that available to typically BiCI users (Exp 2).

**FIG. 1.**
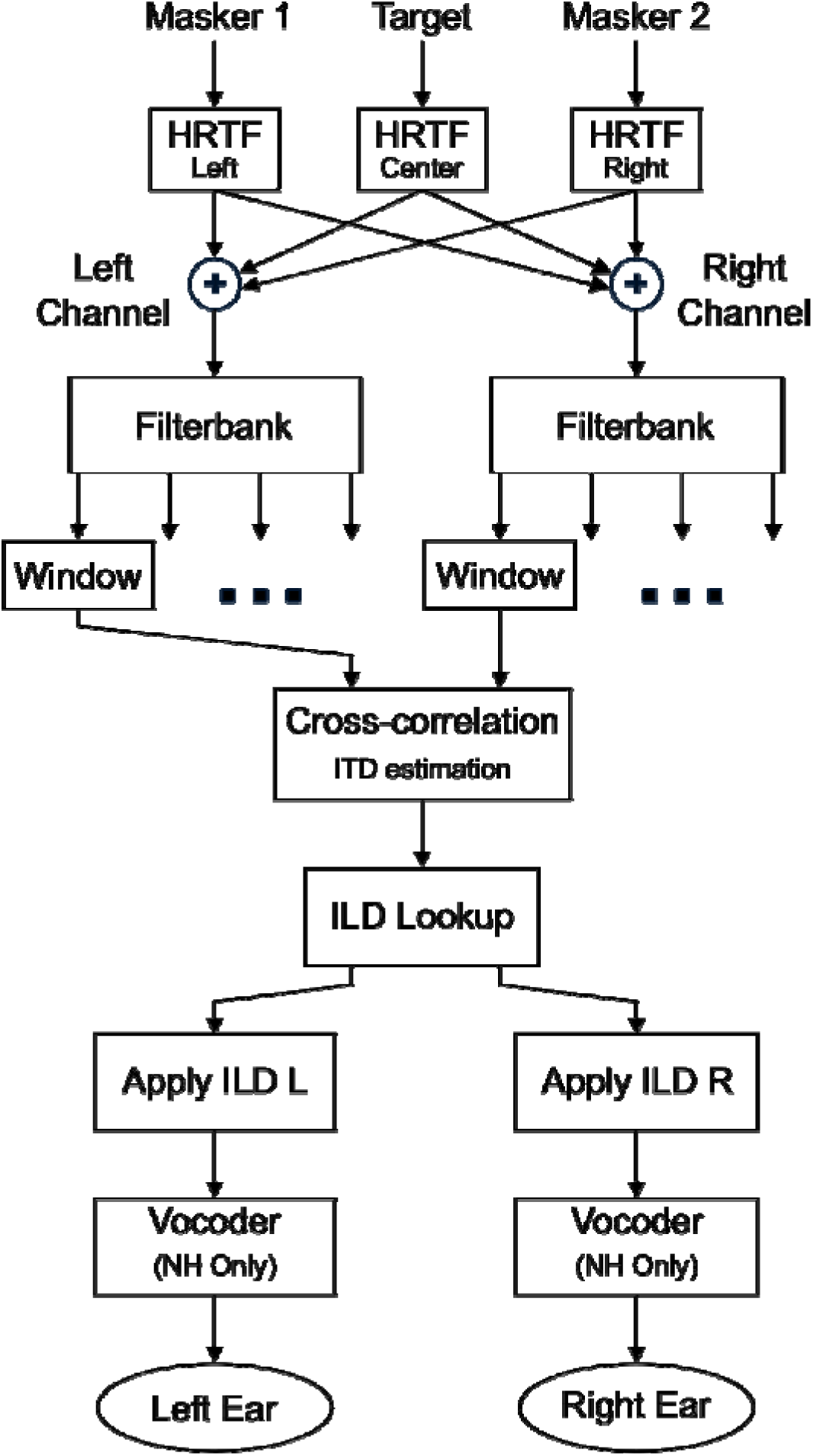
Signal processing flow diagram. Target and masker sound stimuli were first spatialized using HRTFs. Then, working on the combined signal in the left and right ears, signals were filtered into frequency bands, and windowed into 20 ms bins. For each time-frequency bin, the peak of the cross-correlation was used to estimate an ITD. This ITD was converted to an ILD with a lookup table, and applied via contralateral attenuation. Finally, the stimuli were vocoded (in Experiment 1 only).

In the No magnification condition, no further spatial processing was performed (stopped after “Bandpass Filter” in Fig. 1), and vocoding was applied (Exp 1 only). In the ILD magnification conditions, target-masker mixtures were filtered into different frequency bands, with the number depending on the processing condition (“Filterbank” in Fig. 1). We chose the cutoff frequencies to fix the number of frequency bands in which ILD magnification occurred to four. Both magnification filter banks used 4th-order Butterworth filters. In the Low-Frequency magnification condition, the filter bank comprised five contiguous bands with cutoff frequencies of 125, 250, 500, 1000, 2000, and 5500 Hz. ILD magnification was applied to the lowest four bands.

In the Broadband magnification condition, four contiguous bands were created using cutoff frequencies of 125, 300, 900, 3500, and 5500, and ILD magnification was applied to all four bands, matching the number of processing bands in Low-frequency magnification. As described in the Introduction, we were interested in whether fewer, wider bands would provide similar benefit to speech intelligibility as magnification using six independent frequency bands (Brown, 2018).

Following filtering, non-overlapping 20-ms boxcar-shaped windows of data were created within each frequency band (“Window” in Fig. 1). In each window, an ITD was estimated by finding the interaural delay corresponding to the lag of the maximum output of the cross-correlation of the left and right channels (“cross-correlation ITD estimation” in Fig. 1). We defined a maximum ITD of 750 μs, and only cross-correlation maxima in the range [-750,750] were considered. This was to avoid implementing ITDs outside of the natural range that might result from spurious cross-correlation maxima due to interactions across the three sound sources. A lookup table was used to convert the ITDs to ILDs. Specifically, a zero-μs ITD corresponded to a 0-dB ILD, a 750-μs ITD corresponded to a 32 dB ILD, and intermediate values were linearly interpolated, resulting in a linear weighting of approximately 0.043 dB/μs. The resulting ILD in each 20-ms segment was then achieved by attenuating the signal for the ear contralateral to the direction of the ITD for that frequency band and time window, leaving the ipsilateral ear segment unchanged. Although this processing approach causes intensity transitions from window to window that are heard as a light ‘static’ sound when listening to the broadband stimuli, pilot testing indicated that this static is not audible to either BiCI users or NH listeners presented with vocoded stimuli.

It is worth noting that we included the 0-degree configuration (where target and maskers were all co-located) in our experiments as a reference. Although stimuli in the 0-degree configuration went through the same processing pipeline, Low-frequency and Broadband ILD magnification had a negligible effect on the stimuli when target and maskers were co-located at 0-degrees azimuth. Specifically, in this spatial configuration, estimated ITDs were essentially zero in all time-frequency analysis bins; therefore, ILD magnification did not change the signal levels in any substantive way. Moreover, by putting these stimuli through the same processing pipeline, we ensured that any artifacts from the processing were present in this reference condition, as well.

Finally, stimuli for NH listeners (Experiment 1) were processed with an 8-channel sinusoidal vocoder to make the available information similar to what a CI user could access. The first stage of the vocoder was a filterbank with cutoff frequencies spaced logarithmically between 125 and 5500 Hz. In each band, the amplitude envelope was extracted via half-wave rectification and low-pass filtering at the lesser of 400 Hz or half the bandwidth. These envelopes were then used to modulate zero-phase sine tone carriers with frequencies at the logarithmic center of the bands, which were then summed. Because the carriers were zero-phase, they delivered a constant ITD of 0 μs.

### C. Equipment & Procedures

#### 1. Experiment 1 (NH Listeners)

All digital signals (sampling frequency 44.1 kHz) were converted to analog signals with an RME Fireface UFX+ soundcard, and adjusted to an overall presentation level of 70 dB SPL with Atlas AT100-RM passive attenuators. Listeners wore Etymotic ER-3a insert phones. All participants sat in an audiometric booth during testing while the experimenter sat in an acoustically isolated, but physically adjoined booth with a window into the testing room and an intercom allowing verbal communication between participants and the experimenter.

Listeners first heard ten target sentences presented in quiet to familiarize them with the normal acoustic voice of the target talker and then heard 10 vocoded target sentences. Listeners were instructed to verbally repeat back as many of the words as possible produced by the centrally located “target” talker and to guess if they were not sure. Ten sentences, each with an average of five keywords (nouns, verbs, adjectives) for a total of 50 keywords, were presented in each ILD magnification (None, Low-frequency, Broadband) and masker location (± 0, 15, 30, 45, 60, 75, and 90 degrees) condition. No participant ever heard a sentence more than once. For each condition, we quantified performance by calculating the total number of target keywords out of 50 that were correctly identified.

#### 2. Experiment 2 (BiCI Users)

Equipment and procedures were nearly identical to those used in Exp. 1, except for a few minor differences. Specifically, instead of a 70-dB SPL presentation level, BiCI users used the attenuators to adjust a practice trial to a comfortable level. Instead of Etymotic ER-3 insert phones, direct-connect accessory cables connected to the CI speech processors delivered sound. BiCI users used their everyday programs with no adjustments, including for automatic gain control. Finally, BiCI practice consisted of 10 unprocessed target sentences in quiet.

## III. RESULTS

### A. Experiment 1

Figure 2 presents the across-subject average results for Experiment 1 (NH listeners), showing the raw percent correct speech intelligibility for each condition as a function of target-masker separation. In the co-located control condition, listeners correctly reported roughly 50% of the keywords. Although performance in the co-located condition is plotted separately for each ILD magnification, they represent acoustically similar trials since all sound sources were located at midline– the only differences were that the stimuli in the two ILD magnification conditions went through multiple processing steps that introduced modest artifacts. In the No magnification condition, performance was similar to the co-located configuration for less lateral maskers (±15, 30 & 45°), but was better for more lateral masker locations (±60, 75 & 90°). In contrast, for both of the ILD magnification conditions, performance was similar for all spatially separated masker conditions; performance in the separated configurations was better than for the co-located configuration. These results show that ILD magnification improves target speech intelligibility compared to listening with naturally occurring ILDs when the maskers are spatially separated from the target but relatively close to midline.

**FIG. 2.**
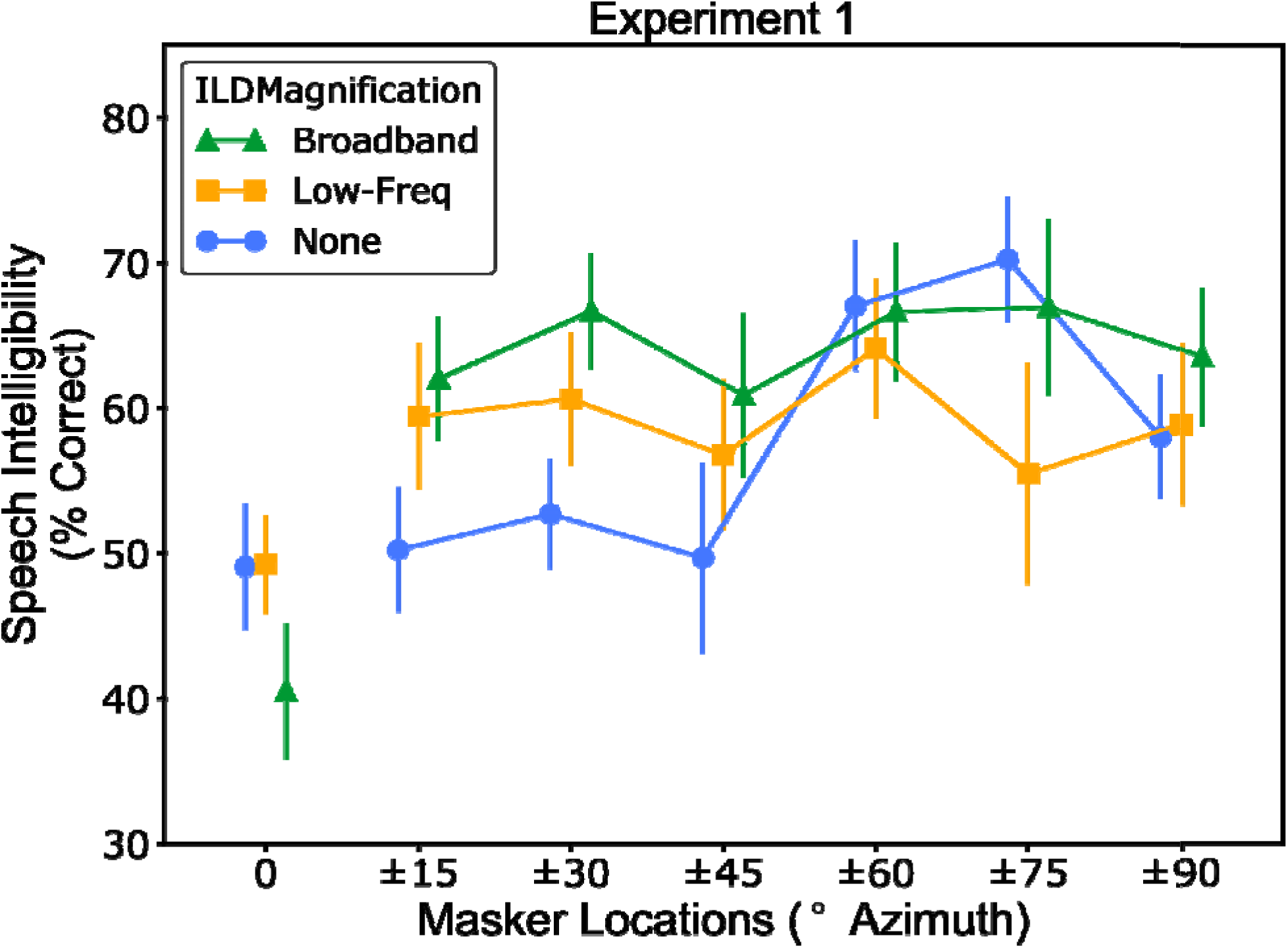
Experiment 1 Results. Average percent keywords correct as a function of masker location for sixteen NH listeners in a BiCI (vocoded) simulation. Data are shown for the three ILD magnification conditions: None (circles), Low-frequency (square), and Broadband (diamonds). The symbols show the across-subject average and the error bars show ±1 across subject standard error.

A Shapiro-Wilk test of normality was conducted to determine whether percent correct data at spatially separated masker locations are normally distributed. The results indicate that the data are normally distributed (p = 0.143). Additionally, Mauchly’s test indicated that in no case was the assumption of sphericity violated (all p > 0.05); accordingly, no sphericity-based corrections were applied to our statistics. A 2-factor repeated-measures ANOVA was computed with Masker location and ILD magnification as the two main factors and percent-correct speech intelligibility as the dependent factor. Significant effects were observed for the main factor of Masker location (F(6,90) = 10.43, p = 9.30e-9, ^2^ = 0.088) and the interaction between ILD magnification and Masker location (F(12,180) = 2.59, p = 3.00e-3, ^2^ = 0.044); the effect of ILD magnification was nearly significant (F(2,30) = 2.89, p = 0.071, ^2^ = 0.009). To interpret the significant interaction, we conducted follow-up one-way ANOVAs separately for each masker location. There was a significant effect of ILD magnification in the ±15 degree (F(2,30) = 3.72, p = 0.036, ^2^ = 0.076) and ±30 degree (F(2,30) = 7.81, p = 0.002, ^2^ = 0.113) conditions. Bonferroni-corrected pairwise t-tests within each masker location showed that performance at ±15° was greater with Low-frequency (p = 0.025) and Broadband (p = 0.047) magnification than with No magnification. At ±30°, both Broadband (p = 0.013) and Low-frequency (p = 0.050) magnification increased percent-correct keyword performance. This pattern of results suggests that ILD magnification improved task performance for NH listeners, but only at small target-masker separations.

### B. Experiment 2

Figure 3 presents the across-subject average results for Exp. 2 in 7 BiCI users, showing the raw percent correct speech intelligibility for each condition as a function of target-masker separation. In this population, performance in the co-located condition (about 40 percentage points) was generally lower than in the spatially separated conditions.

**FIG. 3.**
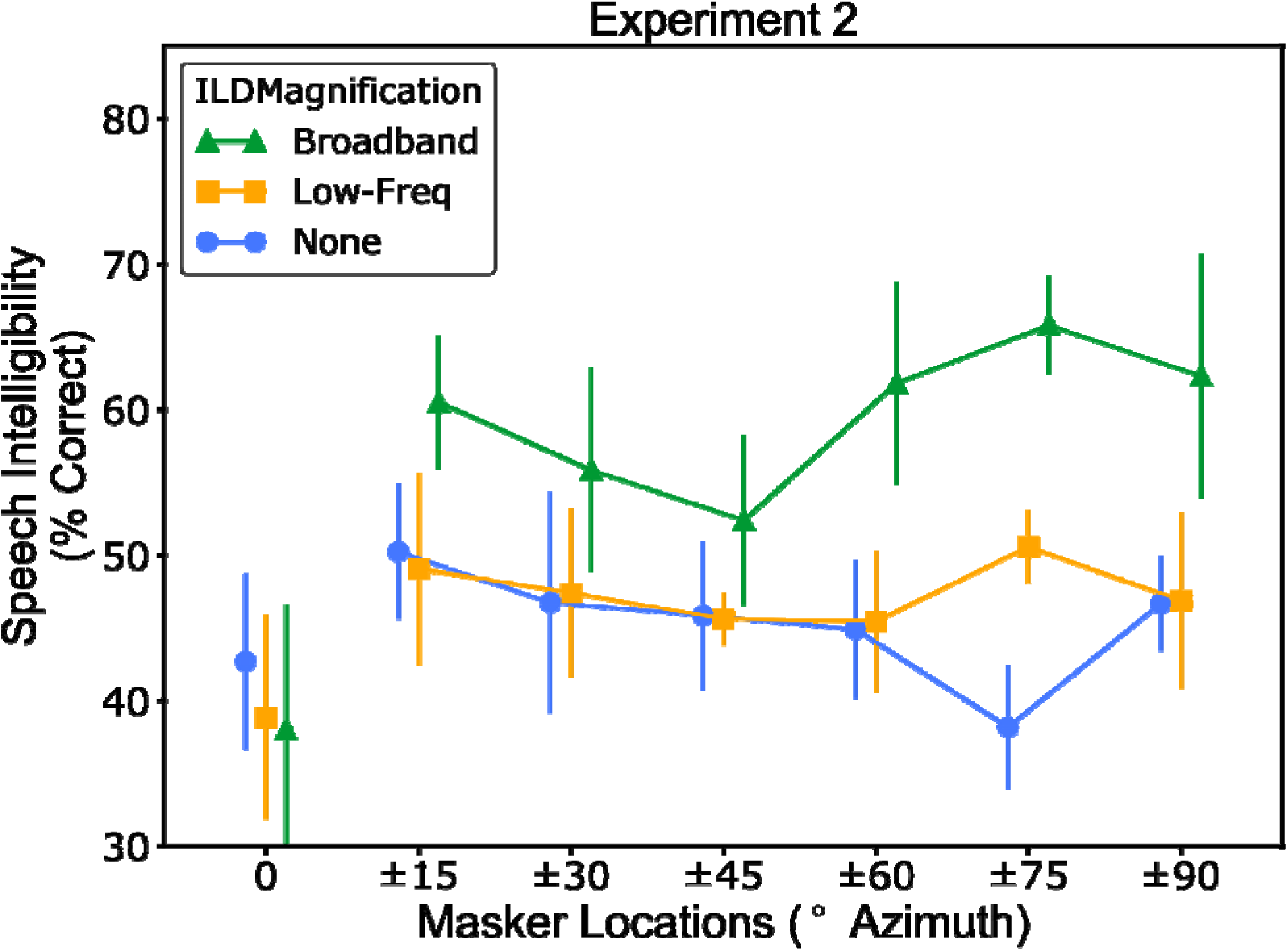
Experiment 2 Results. Average percent keywords correct as a function of masker location for seven BiCI users. Data are shown for the three ILD magnification conditions: None, Low-frequency, and Broadband. The symbols show the across-subject average and the error bars show across-subject ±1 standard error in each condition.

A Shapiro-Wilk test of normality failed to reach significance (p = 0.651), so we treated the data as normally distributed. Sphericity was also not violated in our measurements (Mauchly’s test, all p > 0.05), so no additional sphericity corrections were applied. A 2-factor repeated-measures ANOVA was conducted with main factors of masker location and ILD magnification, and percent-correct speech intelligibility as the dependent variable. The ANOVA revealed significant main effects of ILD magnification (F(2,12) = 11.50, p = .002, ^2^ = 0.092) and Masker location (F(6,36) = 2.65, p = 0.033, ^2^ = 0.069). The interaction between them was not significant (F(12,72) = 0.88, p = 0.57, ^2^ = 0.039). Bonferroni-corrected post hoc two-tailed paired t-tests revealed that the Broadband ILD magnification strategy produced performance that was significantly better than both No magnification (p < 0.001) and Low-Frequency magnification (p < 0.001). However, there was no significant difference between performance in the No magnification and Low-Frequency magnification conditions (p = 0.59). This pattern of data suggests that Broadband ILD magnification improved performance for BiCI users. Unlike NH listeners, however, this effect was observed at all spatial separations rather than only small spatial separations (i.e. we did not observe an interaction). We also conducted post-hoc tests across masker location. These showed that performance in the co-located condition was significantly poorer than ±15° (p = 0.02) and ±90° (p = 0.038), although the differences in percent correct are small. Performance did not vary between any spatially separated masker location conditions.

## IV. DISCUSSION

### A. ILD Magnification improved speech intelligibility through a binaural mechanism

Broadband ILD magnification significantly improved speech intelligibility in a symmetrical-masker task for both NH listeners hearing vocoded stimuli (Experiment 1) and BiCI users (Experiment 2). Low-frequency magnification, however, only had a significant effect for NH listeners at small target-masker separations. Given that our ILD manipulations should not appreciably change monaural TMR (see below), the benefit of ILD magnification must be binaural in nature. As we discuss below, ILD magnification may cause modest changes in loudness, but the largest possible loudness changes are too small to explain our results. We thus argue that ILD magnification increases the perceived spatial separation between the target and the more salient masker in the time instances and frequencies where one masker dominates the sound mixture. To support this explanation, we first analyze the consequences of ILD magnification to rule out simple acoustic explanations for the effects we observed.

With naturally occurring ILDs, the better-ear effect contributes to improved speech intelligibility, and this effect grows as the separation between target and masker increases. One might reasonably wonder, then, whether introducing ILD magnification simply increases better-ear effects (i.e., increases monaural TMR), rather than increasing perceived target-masker separation. Non-adaptive beamformers, for example, facilitate SRM through both changes in monaural TMR and ILD-based localization cues (Yun et al., 2021). However, the ILD magnification algorithms presented here work by attenuating the summed target-masker mixture at one ear, as dictated by the estimated ITD in a given spectro-temporal bin. That is, both the target and masker in a given ear are attenuated by the same amount. Thus, the acoustic TMR at either ear is not altered, logically ruling out any effects due to an increased better-ear benefit.

We confirmed that monaural better-ear effects were unchanged in our stimuli and that binaural loudness effects could not explain improvements in performance by analyzing the final stimuli for spatial configurations used in this study after applying our processing. Specifically, we spatialized and summed stimuli as in the experiments, bandpass filtered and time windowed the left and right channels, and calculated the estimated ITDs and associated ILD adjustments (the “gain tracks”) to be applied at each time and in each frequency in each ILD magnification condition and for each masker azimuth.

An example of these gain tracks is shown in Fig. 4. Panel A depicts the symmetrical-masker spatial configuration, with example waveforms for each source. Panel B plots the applied ILDs in each time-frequency bin for the Low-frequency (upper panel) and Broadband (lower panel) magnification conditions. The particular example in the figure placed maskers at ±60°. The waveforms in Panel A are time-aligned to the ILD tracks in Panel B. The legend at the bottom of the figure shows how ILD magnitudes correspond to the different colors in Panel B. The highest frequency band in the Low-frequency magnification plot (2 – 5.5 kHz) is solid blue, because ILD magnification was not applied to this band. It is visually apparent that when a given sound source is more intense than the others (dominates) in the mixture, it drives the ITD estimate at that moment in time (for example, Masker 1 drives the ITD estimation at around 0.25 s, whereas the Target drives it at around 1.1 s).

**FIG. 4.**
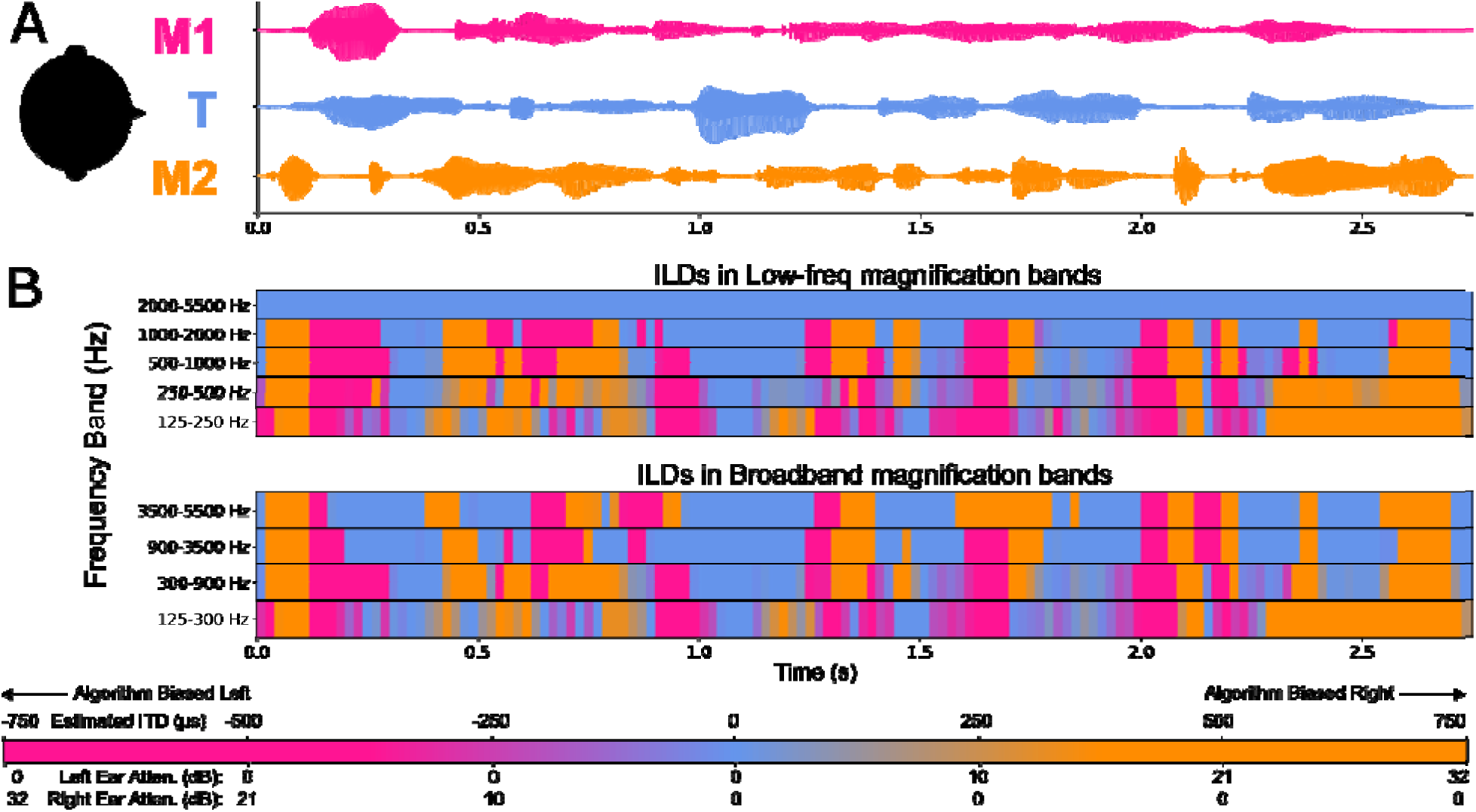
Acoustic analysis example sentence. Panel A shows the relative positions of the target (T), masker 1 (M1) and masker 2 (M2), along with corresponding example waveforms. Panel B shows applied ILDs in spectro-temporal bins for Low-frequency (top plot) and Broadband (bottom plot) magnification. In this example, maskers were spatialized to +/-60 degrees. The legend at the bottom maps color to the applied attenuation in each ear (i.e., the ILD), and the estimated ITD for reference. The waveforms in Panel A and the ILD plots in Panel B are time-aligned, making it easier to observe that when a given sound source is more intense than the others, that source biases the ITD estimation toward its azimuth. Eg., when M1 (spatialized to the left) is relatively high at around 0.25 s, the ILD applied is to the left. Note that the highest frequency band in Low-frequency magnification (2000-5500 Hz) always shows a 0 dB ILD, as no magnification was applied in this band.

We calculated the TMR in each time-frequency window with and without ILD magnification by applying the ILD gain tracks to the unmixed target and masker signals independently (actual ILD magnification always works on the mixed signals). We found that the difference in TMR between naturally occurring ILD stimuli and ILD magnified stimuli at any given time-frequency bin for any stimulus was on the order of a rounding error (ie., less than about 1e-13), as expected (given that the same gain track was applied to the target, dominant masker, and weaker masker). This was true for all masker azimuths: regardless of the magnitude of the level change in a given time-frequency bin, the target and maskers were all equally attenuated in the contralateral ear, resulting in no change in monaural TMR. Thus, changes in monaural TMR cannot explain our results.

As shown in Figure 4, our ILD magnification approach only changes signals in time instances and frequency bands where one of the maskers dominates. In those time-frequency bins, the algorithm reduces the sound level of the mixed signal reaching the ear contralateral to the dominant masker. This can cause reductions in the perceived loudness of the target as well as both masker sounds, which could contribute to changes in speech intelligibility results. However, studies of binaural loudness summation show that reductions in binaural loudness are considerably less than would be predicted strictly from the acoustics (Epstein and Florentine, 2009). For instance, going from a diotic presentation to a monaural presentation reduces the total sound energy summed across both ears by 3 dB. Yet, the perceived change in binaural loudness for a diotic signal vs a monotic signal is considerably less than changing the monaural level by 3 dB. For a sound that is more intense in one ear, reducing the signal level of the ear with the less intense sound will change binaural loudness even less than for a diotic sound. The only large change in binaural loudness caused by our ILD magnification approach would affect the non-dominant masker, where we attenuate the ear signal that contains the more intense sound. In such instances, the binaural loudness will depend on the levels of the signals reaching each ear and the amount of attenuation applied. In the most extreme case, if the sound level in the less intense ear was too quiet to be detected and the attenuation at the more intense ear dropped the sound level in that ear below detection threshold, attenuation could render that signal inaudible.

In all of our tested spatial configurations, the target is located at midline; therefore, ILD magnification will reduce the binaural loudness of the target by 3 dB at most. Similarly, the dominant masker source loudness will decrease by less than 3 dB– and typically by even less than the diotic target loudness changes, since in most frequency bands, the dominant masker will already be less intense in the ear that is attenuated. Thus, our algorithm will either cause a decrease in or leave unchanged the target-to-dominant-masker loudness ratio, since the dominant masker loudness cannot be decreased by more than that of the target loudness.

It may be that the non-dominant masker’s binaural loudness is reduced by the algorithm by more than any changes in the binaural loudness of the target, which could contribute to increased intelligibility. In the most extreme case, the non-dominant masker may change from being audible to being inaudible due to magnification. However, practically speaking, the non-dominant masker is unlikely to contribute to masking in the time-frequency bins where our “gain track” calculation causes a large attenuation, since the non-dominant masker is, on average, less intense than the dominant masker in those moments and frequencies.

This is demonstrated in Figure 5A, which plots the mean (and standard error) of the energy ratio of the left masker vs. the right masker as a function of the calculated ITD (and thus the applied ILD), averaged across individual time-frequency analysis windows. The plot only includes time-frequency bins where the target does not dominate (i.e., this analysis excludes time-frequency voxels when the target is more intense than each of the maskers; the applied ILD in those cases is almost always close to zero). The maskers were placed at ±90° for this analysis. In those time-frequency bins in which attenuation is applied, the non-dominant masker intensity typically is already much less intense than the dominant masker – almost never 3 dB or less, most often at least 10 dB, and on the order of 20 dB when the applied attenuation is greatest – strongly suggesting that attenuation from magnification is unlikely to contribute significantly to reduced loudness of the non-dominant masker relative to the target.

**Figure 5.**
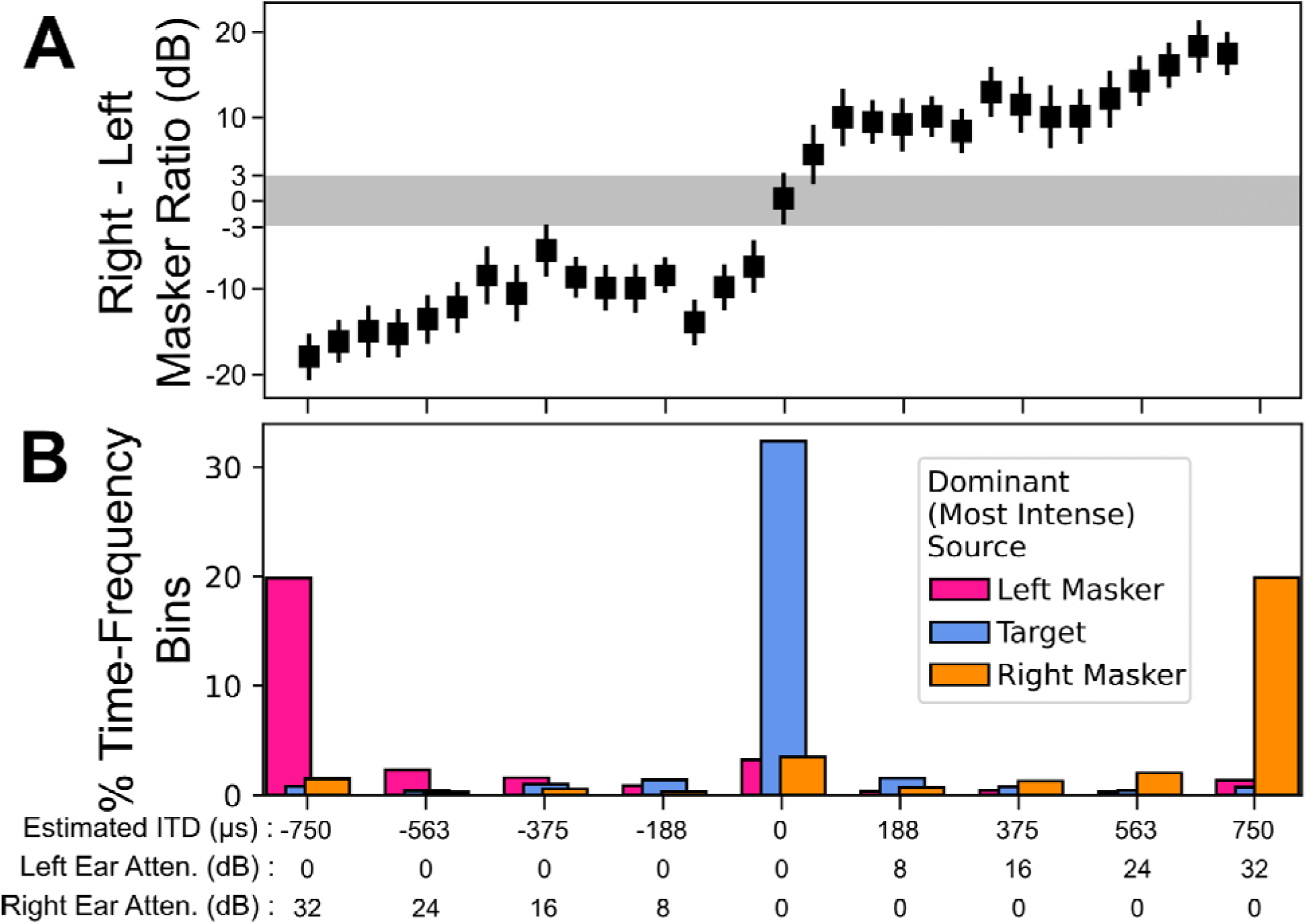
Source-to-source ratio analysis. Panel A shows the mean and standard error of the energy ratio between right and left masker sounds over time-frequency bins in 20 example stimulus sets as a function of the estimated ITD (and therefore the to-be-applied attenuation in each ear). Only time-frequency bins in which the target does not dominate are shown: bins in which the target was greater than 3 dB more intense than the combined masker signals are excluded. The estimated ITD shown is for maskers placed at ±90°. The gray shaded error shows the limits of ±3 dB right to left masker ratio: all points where non-zero attenuation is applied are outside this region. Panel B shows the percentage of time-frequency bins in which each sound source dominates as a function of estimated ITD. Values are binned based on equally spaced estimated ITD values (±750 μs, ±563 μs, ±375 μs, ±188 μs,0 μs), and bin widths of 188 μs. Large attenuation is only applied when a given source is more intense than the others. Therefore, a sound source will not be attenuated when it dominates the mixture and drives perceptual masking.

Figure 5B illustrates that the algorithm enhances the perceptual separation between sound sources. Specifically, the distribution of calculated ITDs across time-frequency bins has a pronounced trimodal distribution, with peaks that correspond to far left, center, and far right; these modes in turn correspond to cases where the most intense source in a given time-frequency bin is the left masker, center target, or right masker, respectively. This analysis demonstrates that, e.g., left-indicating ILDs are applied by the algorithm when the left source dominates the mixture. Together, Figures 5A and 5B support the argument that the main effect of ILD magnification is to enhance the perceived laterality of lateral sources that dominate the sound mixture, rather than by reducing the binaural loudness of the nondominant masker.

In BiCI users, insertion depths, neural survival, and programming differences may lead to changes in binaural fusion. Binaural fusion in BiCI users is strong for pitch and often spans a wide range (Reiss et al., 2014, 2018). In our stimuli, if BiCI users show strong fusion, this may lead to poor segregation. This is something that the perceptually salient ILDs we create addresses. If different bands are dominated by different sources, they may be less likely to fuse and segregation will improve because of the imposed ILDs causing greater perceptual separation in a given frequency band. It may also be that binaural fusion is reduced due to interaural asymmetries in array insertion depth (Kan et al., 2013). If a BiCI user is receiving two slightly different, unfused versions of the scene, then contributions of the dominant masker in the to-be-attenuated ear may add interference, but not produce perceived spatial separation between target and masker because of the reduced fusion. In this case, attenuation from the applied ILD may simply be reducing the intensity of what is essentially a monaural interferer– the dominant masker in the contralateral ear. However, binaural fusion appears to be, if anything, overly broad for BiCI users (Reiss et al., 2014, 2018). Thus, a lack of binaural fusion, particularly for broadband stimuli like speech, does not seem to be a likely contributor to our results. In addition, poor fusion would not explain the performance of the NH group, who should have good fusion in any event. Given that the patterns of results across the two experiments track closely with one another and appear to be binaural in nature, this possibility seems unlikely to be a significant factor in the outcome of the current study.

In NH listeners in BiCI simulation, we hypothesized that both Low-frequency and Broadband magnification would significantly improve performance compared to No magnification (ie., naturally occurring ILD cues). Supporting these hypotheses, we observed a significant benefit from Low-frequency and Broadband magnification, but only at small spatial separations (±15° and ±30°). These results are generally in line with a previous study that showed a significant increase in intelligibility even for low-frequency ILD magnification using different ILD magnification algorithm parameters and a different spatial configuration (Brown, 2014). The spatial configuration used here, with a masker to either side of the target, poses much greater challenges, both for the algorithm and for the listener, than when hearing a target to one side and a single masker to the other. Better-ear acoustic effects only occur in glimpses (moments and frequencies where one of the two maskers happens to be low in overall intensity) in the symmetric masker configuration. Together, these differences produce much more perceptual interference in the symmetrical masker configuration than the spatial configurations of this previous study, which may explain why there is no significant effect of Low-frequency ILD magnification in that previous work. The more challenging listening situation tested in the current study may require spatial separation in higher frequency regions important for speech intelligibility (DePaolis et al., 1996; Hogan and Turner, 1998; Vickers et al., 2001). Consistent with this hypothesis, Broadband magnification yielded significantly greater benefit than Low-frequency magnification.

In BiCI users, we hypothesized that both Low-frequency and Broadband magnification would produce greater benefit from spatial separation between target and maskers than naturally occurring ILD cues. We observed a significant improvement in performance with Broadband magnification over No magnification; however, Low-frequency magnification provided no statistically significant benefit. This effect did not interact with masker location; that is to say, Broadband magnification improved speech intelligibility for all masker locations. It is worth noting that we observed significant effects of Broadband magnification here even though the subject pool comprised only 7 listeners. The lack of statistically significant differences between the No magnification and Low-frequency magnification may be due to a lack of power rather than a failure of ILD magnification. To estimate how many participants would be needed to observe an effect of Low-frequency magnification, we conducted a post-hoc power analysis using G*Power (Faul et al., 2007). This analysis suggested that 32 BiCI users would be needed to observe a difference between No magnification and Low-freq magnification with power of 0.80 and α = 0.05.

For BiCI users, naturally occurring ILD cues failed to produce SRM, even with maskers at ±90°. Only in the magnified ILD conditions did spatial separation yield performance that was better than in the co-located configuration. This result supports our hypothesis that BiCI users require greater ILDs to perceive a separation between target and maskers. This is also consistent with the literature (Brown, 2014; Ihlefeld and Litovsky, 2012), and is likely due to the compressed perceptual space BiCI listeners experience for sources in the horizontal plane with naturally occurring cues (Grantham et al., 2007). ILD magnification can mitigate this limitation for BiCI users by expanding the perceptual space (Brown, 2018). While neither naturally occurring ILDs (No magnification) nor Low-frequency ILD magnification improved performance for our BiCI users compared to the co-located configurations, Broadband ILD magnification did. Given that our ILD magnification approach does not alter the TMR at either ear, these results suggest that Broadband ILD magnification allows BiCI listeners to more effectively deploy spatial selective attention, focusing on the target stream and suppressing the maskers. However, neither naturally occurring ILDs nor Low-frequency ILD magnification allowed the BiCI users to focus spatial attention effectively.

Though we demonstrate that ILD magnification improved speech intelligibility in both NH listeners hearing vocoded signals and in BiCI users, there are a number of questions our study cannot answer. For instance, vocoding only captures first-order effects of listening to BiCI signals (ignoring current spread, mismatches in binaural stimulation, compressed level dynamic range, etc.). Thus, it may not be surprising that our two groups showed different patterns of results. Our two groups also differed in age demographics (NH listeners, 19-37 years; BiCI users, 52-76 years). Future studies could address how such factors influence the utility of ILD magnification.

The overall benefit of ILD magnification observed in the current study is less than what was observed when listeners heard a target presented with a single masker (Brown, 2014). In this earlier non-symmetric masker study, ILD magnification increased percent correct performance by about 30 percentage points. In contrast, Broadband magnification in the current study improved performance by around 20 percentage points over no magnification. This difference likely reflects the perceptual difficulty associated with the symmetrical masker configuration used here. Another potential factor is the number of processing bands, which was reduced from 20 in the previous study to 4 here. Follow-up studies are needed to establish the relationship between processing band number and speech benefit. Nevertheless, 20 percentage points of masking release represents a substantial benefit and indicates that ILD magnification can be effective in relatively complex auditory scenes.

The question remains as to the mechanism by which ILD magnification provides speech intelligibility benefit. As we argue above, we believe that any changes in TMR or in binaural loudness caused by ILD magnification are too modest to account for the improvements we observe. Instead we propose that magnified ILDs enhance perceived spatial separation between sound sources, which enhances binaural unmasking (Freyman et al., 1999). This enhanced spatial separation allows spatial selective attention to be deployed more successfully. NH listeners can use auditory spatial cues to selectively attend to a target amongst spatially separated maskers (Noyce et al., 2021; Shinn-Cunningham, 2017; Shinn-Cunningham and Best, 2008). This process occurs through coordinated activity in multiple areas of the brain, including prefrontal cortex, parietal cortex, and auditory cortex (Alho et al., 2014; Choi et al., 2014; Deng et al., 2019a, 2019b; Noyce et al., 2022). Spatial auditory attention is less effective in listeners with hearing loss; indeed, performance is inversely correlated with spatial discrimination thresholds (Bonacci et al., 2019; Dai et al., 2018). Similarly, limitations of electrical stimulation reduce CI users’ ability to capitalize on naturally occurring spatial cues to direct selective attention (Akbarzadeh et al., 2020; Goupell et al., 2016). This underscores the need for processing algorithms like ILD magnification that facilitate greater perceptual segregation and more effective deployment of spatial attention.

### B. Corrective ILD Magnification and Future Work

The current approach maximizes the perceptual separation between target and masker, which provides significant improvement in speech intelligibility. But there are other spatial configurations in which the current strategy will likely be less effective, if not detrimental. Specifically, if a target and masker are spatially separated, but on the same side of midline, ILD magnification may decrease the perceptual separation between them. This is because there is a limit to the perceived lateral position of a sound source. Aggressive ILD magnification like that used in the current study may cause ipsilateral sources to be hyper-lateralized (sources perceived to be as far to the side as possible), which may actually lead to reduced perceptual separation between them.

Corrective ILD magnification (Brown, 2018) may represent a potential compromise. Whereas the ILD magnification strategy used in the current study was designed to maximize the perceptual separation between target and maskers for our configurations (with the target at midline and maskers to the sides), corrective processing is designed to minimize rms error between perceived and actual locations of the sources in the mixture. When tailored to individual BiCI users, corrective ILD magnification significantly improves localization accuracy; two users presented with this strategy exhibited localization performance on par with a group of NH listeners. Individually tailored corrective ILD magnification could also address confounds of automatic gain control or electrode inactivation, which we did not control for in this study since BiCI users used their everyday programs. Future studies will likely need to strike a balance between maximizing SRM task performance, as in the current study, and maximizing perceptual accuracy of source location (Brown, 2018).

Continued research should further characterize the frequency-, azimuth-, and subject-specific benefits of magnified ILD cues in an SRM paradigm. Important parameters to explore include the number of magnification bands, the cutoff frequencies of those bands, and the ITD-to-ILD mapping function (lookup table). Four magnification bands were used in this study to explore whether SRM benefits could be obtained using a small number of frequency bands, which is less computationally demanding than similar past approaches (Brown, 2014). Relatedly, the optimal processing bandwidth will also need to be established. Both the number of bands and the bandwidths that provide the maximum benefit may be frequency-, azimuth-, or subject specific.

The algorithm we used in the current study was specifically designed to manipulate ILDs without requiring any a priori knowledge of the specific locations of sound sources; the algorithm will accentuate the spatial separation between a central target and maskers to one or both sides of the listener, or when the target is on one side of the head and any maskers are in the opposite hemifield. This allows the algorithm to be effective even in relatively complex acoustic environments like the symmetrical-masker configuration employed in the current study. It also works best with modulated maskers (as it operates during “glimpses” where one masker dominates), which have proven to be more difficult for traditional noise reduction approaches. But there are spatial configurations that may pose a problem. For example, if a target and masker are spatially separated but on the same side of midline, ILD magnification may actually reduce the perceptual separation between them. Future work should explore this possibility.

We argue that the observed benefits for understanding target speech come from enhancing the perceived spatial separation between target and maskers. Future experiments can explicitly examine this hypothesis by measuring neural responses to sound in both normal hearing listeners and BiCI users completing spatial selective attention tasks with and without ILD magnification. If the magnified ILD cues used here do, in fact, allow for greater sound source segregation, this should be evident from neural signatures of spatial selective attention.

## V. ACKNOWLEDGEMENTS

This work was supported in part by NIH grants R21-DC018408 to CAB and R01-DC015988 to BGSC. We would like to thank Sydney Sepkovic and Cece Restauri for help with data collection.

## VII. ETHICS STATEMENT

This study was reviewed and approved by The University of Pittsburgh Institutional Review Board. The patients/participants provided their written informed consent to participate prior to their participation in this study.

## VIII. DATA AVAILABILITY STATEMENT

The raw data supporting the conclusions of this article will be made available by the authors, without undue reservation.

